# Paradoxical phase response of gamma rhythms facilitates their entrainment in heterogeneous networks

**DOI:** 10.1101/2020.12.15.422838

**Authors:** Xize Xu, Hermann Riecke

## Abstract

The synchronization of different *γ*-rhythms arising in different brain areas has been implicated in various cognitive functions. Here, we focus on the effect of the ubiquitous neuronal heterogeneity on the synchronization of PING (pyramidal-interneuronal network gamma) and ING (interneuronal network gamma) rhythms. The synchronization properties of rhythms depends on the response of their collective phase to external input. We therefore determined the macroscopic phase-response curve for finite-amplitude perturbations (fmPRC), using numerical simulation of all-to-all coupled networks of integrate-and-fire (IF) neurons exhibiting either PING or ING rhythms. We show that the intrinsic neuronal heterogeneity can qualitatively modify the fmPRC. While the phase-response curve for the individual IF-neurons is strictly positive (type I), the fmPRC can be biphasic and exhibit both signs (type II). Thus, for PING rhythms, an external excitation to the excitatory cells can, in fact, delay the collective oscillation of the network, even though the same excitation would lead to an advance when applied to uncoupled neurons. This paradoxical delay arises when the external excitation modifies the internal dynamics of the network by causing additional spikes of inhibitory neurons, whose delaying within-network inhibition outweighs the immediate advance caused by the external excitation. These results explain how intrinsic heterogeneity allows the PING rhythm to become synchronized with a periodic forcing or another PING rhythm for a wider range in the mismatch of their frequencies. We demonstrate a similar mechanism for the synchronization of ING rhythms. Our results identify a potential function of neuronal heterogeneity in the synchronization of coupled *γ*-rhythms, which may play a role in neural information transfer via communication through coherence.

**Author Summary:** The interaction of a large number of oscillating units can lead to the emergence of a collective, macroscopic oscillation in which many units oscillate in near-unison or near-synchrony. This has been exploited technologically, e.g., to combine many coherently interacting, individual lasers to form a single powerful laser. Collective oscillations are also important in biology. For instance, the circadian rhythm of animals is controlled by the near-synchronous dynamics of a large number of individually oscillating cells. In animals and humans brain rhythms reflect the coherent dynamics of a large number of neurons and are surmised to play an important role in the communication between different brain areas. To be functionally relevant, these rhythms have to respond to external inputs and have to be able to synchronize with each other. We show that the ubiquitous heterogeneity in the properties of the individual neurons in a network can contribute to that ability. It can allow the external inputs to modify the internal network dynamics such that the network can follow these inputs over a wider range of frequencies. Paradoxically, while an external perturbation may delay individual neurons, their ensuing within-network interaction can overcompensate this delay, leading to an overall advance of the rhythm.

## 1 Introduction

Collective oscillations or rhythms representing the coherent dynamics of a large number of coupled oscillators play a significant role in many systems. In the technological realm they range from laser arrays and Josephson junctions to micromechanical oscillators [1, 2]. Among the important biological examples are the heart rhythm, the circadian rhythm generated by the suprachiasmatic nucleus [3], the segmentation clock controlling the somite formation during development [4], and brain waves [5]. One prominent brain rhythm is the widely observed *γ*-rhythm with frequencies in the range 30-100Hz. The coherent spiking of the neurons underlying this rhythm likely enhances the downstream impact of the neurons participating in the rhythm. The rhythmic alternation of low and high activity has been suggested to play a significant role in the communication between different brain areas [6, 7]. That communication has also been proposed to be controled by the coherence of the rhythms in the participating brain areas [8–13].

For collective oscillations or rhythms to play a constructive role in a system they need to respond adequately to external perturbations and stimuli. For instance, for the circadian rhythm it is essential that it can be reliably entrained by light and phase-lock to its daily variation. Similarly, if rhythms are to play a significant role in the communication between different brain areas, their response to input from other areas represents a significant determinant of their function. Moreover, the stimulation and entrainment of *γ*-rhythms by periodic sensory input is being considered as a therapeutic approach for some neurodegenerative diseases [14].

Even small perturbations can affect oscillations significantly in that they can advance or delay the oscillations, i.e. they can change the phase of the oscillators. This change typically depends not only on the strength of the perturbation but, importantly, also on the timing of the perturbations and is expressed in terms of the phase response curve (PRC), which has been studied extensively for individual oscillators [15]. For infinitesimal perturbations the PRC can be determined elegantly using the adjoint method [16].

If the collective oscillation of a network of interacting oscillators is sufficiently coherent, that system can be thought of as a single effective oscillator. Consequently, the response of the macroscopic phase of the collective oscillation to external perturbations and the mutual interaction of multiple collective oscillations is of interest. The macroscopic phase-response curve (mPRC) has been obtained in various configurations, including noise-less heterogeneous phase oscillators [17, 18], noisy identical phase oscillators [19, 20], noisy excitable elements [21], and noisy oscillators described by the theta-model [22], which is equivalent to the quadratic integrate-fire model for spiking neurons. Recent work has used the reduction of networks of quadratic integrate-fire neurons to two coupled differential equations for the firing rate and the mean voltage [23], which is related to the Ott-Antonsen theory [24, 25], to develop a method to obtain the infinitesimal macroscopic PRC (imPRC) for excitatory-inhibitory spiking networks [26, 27].

A key difference between the response of an individual oscillator to a perturbation and that of a collective oscillation is the fact that the degree of synchrony of the collective oscillation can change as a result of the perturbation, reflecting a change in the relations between the individual oscillators. Thus, the phase response of a collective oscillation to a brief perturbation consists not only of the immediate change in the phases of the individual oscillators caused by the perturbation, but includes also a change in the collective phase that can result from the subsequent convergence back to the phase relationship between the oscillators corresponding to the synchronized state, which is likely to have been changed by the perturbation [18]. Interestingly, it has been observed that the infinitesimal macroscopic phase response can be qualitatively different from the phase response of the individual elements. Thus, even if the individual oscillators have a type-I PRC, i.e. a PRC that is strictly positive or negative, the mPRC of the collective oscillation can be of type II, i.e. it can exhibit a sign change as a function of the phase [21, 22, 28].

Here we investigate the interplay between external perturbations and the internal interactions among neurons in inhibitory and in excitatory-inhibitory networks exhibiting *γ*-rhythms of the ING- and of the PING-type. We focus on networks comprised of neurons that are not identical, leading to a spread in their individual phases and a reduction in the degree of their synchrony. How does this phase dispersion affect the response of the macroscopic phase of the rhythm to perturbations? Does it modify the ability of the network to follow a periodic perturbation ?

We show that the dispersion in the phase together with the within-network interactions among the neurons can be the cause of a paradoxical phase response: an external perturbation that *delays* each individual neuron can *advance* the macroscopic rhythm. We identify the following mechanism underlying this paradoxical response: external perturbations that delay individual neurons sufficiently allow the within-network inhibition generated by early-spiking neurons to suppress the spiking of less excited neurons. This results in a reduced within-network inhibition, which reduces the time to the next spike volley, speeding up the rhythm. This paradoxical phase response increases with the neuronal hetero-geneity and allows the network to phase-lock to periodic external perturbations over a wider range of detuning. Thus, the desynchronization within the network enhances its synchronizabilty with other networks. The mechanism is closely related to that underlying the enhancement of synchronization of collective oscillations by uncorrelated noise [29] and the enhanced entrainment of the rhythm of a homogeneous network to periodic input if that input exhibits phase dispersion across the network [30,31]. We demonstrate and analyze these behaviors for networks of inhibitory neurons (ING-rhythm) and for networks comprised of excitatory and inhibitory neurons (PING-rhythm).

## 2 Results

We investigated the impact of neuronal heterogeneity on the response of the phase of *γ*-rhythms to brief external perturbations and the resulting ability of rhythms to synchronize to periodic input. As described in the Methods, we used networks comprised of minimal integrate-fire neurons that interact with each other through synaptic pulses modeled via delayed double-exponentials. To study ING-rhythms all neurons were inhibitory, while for the PING-rhythms we used excitatory-inhibitory networks. In both cases, the coupling within each population was all-to-all. Throughout, we implemented the neuronal heterogeneity by injecting a different steady bias current *I*_*bias*_ into each neuron. Our analysis suggests that the origin of the neuronal heterogeneity plays only a minor role as long as it leads to a dispersion of their spike times [29].

### Paradoxical Phase Response of Heterogeneous Networks: ING-Rhythm

In the absence of external perturbations the all-to-all inhibition among the neurons lead to rhythmic firing of the neurons. Due to their heterogeneity they did not spike synchronously but sequentially, as shown in Fig.1A, where the neurons are ordered by the strength of their bias current. The dependence of the phase dispersion on the coefficient of variation of the heterogeneity in the bias current (CV) is shown in Suppl. Figure S1. For sufficiently large heterogeneity some neurons never spiked: while the weak bias current they received would have been sufficient to induce a spike eventually, the strong inhibition that was generated by the neurons spiking earlier in the cycle suppressed those late spikes. Neurons with strong bias current could spike multiple times.

**Figure 1:**
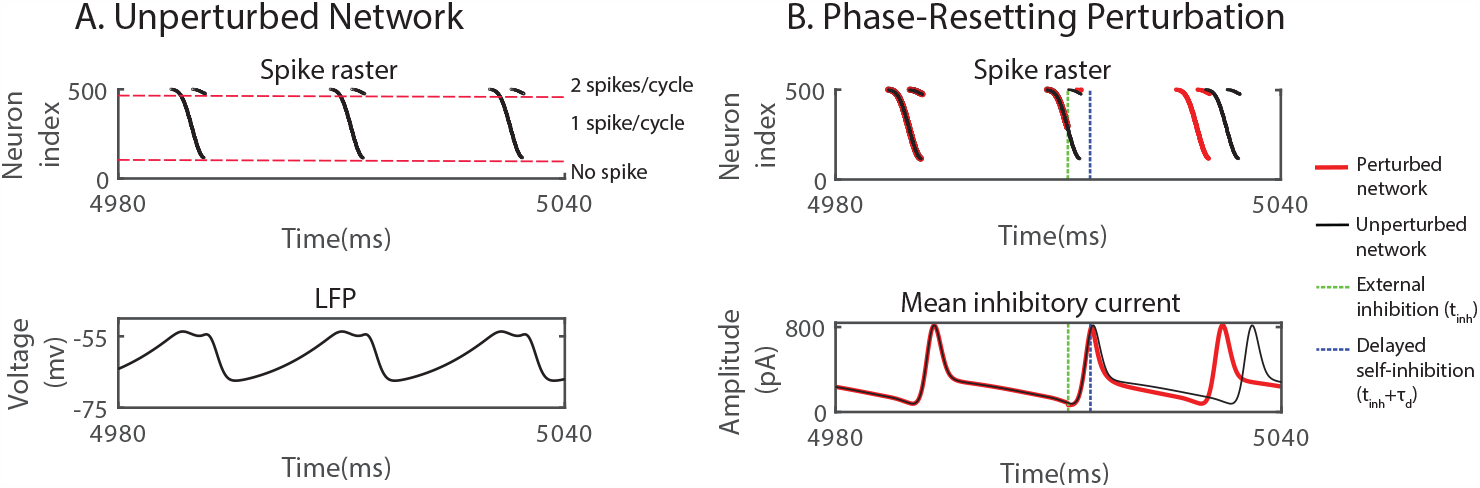
ING-rhythm can be advanced by inhibition while individual neurons are delayed. (A) Top: spike raster of neurons spiking sequentially in the order of their input strength (increasing with neuron index). Bottom: mean voltage across the network (LFP). (B) External inhibition advanced the rhythm. Top: raster plot of spikes without (black) and with (red) external inhibitory pulse. Bottom: Average of the total inhibitory current each neuron received from the other neurons within the network. 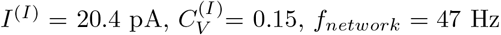. In (B), perturbations were made with a square-wave inhibitory current pulse with duration 0.1 ms and amplitude 3200 pA to each neuron, resulting in a 4 mV rapid hyperpolarization.

A brief, inhibitory external input delivered to all neurons (green dashed line in Fig.1B) delayed each neuron. The degree of this individual delay depended on the timing of the input, as is reflected in the PRC of the individual neurons. If the perturbation was applied during the time between the spike volleys, the delay of each neuron had no further consequence and the overall rhythm was delayed. However, if the same inhibitory perturbation arrived during a spike volley (dashed green line in Fig.1B), it could advance the overall rhythm. As illustrated in Fig.1B, only the spiking of the late neurons was delayed by the perturbation. Importantly, with this delay some neurons did not spike before the within-network inhibition triggered by the early-spiking neurons (dashed blue line in Fig.1B) became strong enough to suppress the spiking of the late neurons altogether. With fewer neurons spiking, the all-to-all inhibition within the network was reduced, allowing all neurons to recover earlier, which lead to a shorter time to the next spike volley. If the speed-up was larger than the immediate delay induced by the external inhibition, the overall phase of the rhythm was advanced by the delaying inhibition.

As the example in Fig.1B shows, the paradoxical phase response requires proper timing of the perturbation. We therefore determined quantitatively the macroscopic phase-response curve (PRC) of the rhythm. To do so we measured computationally the amount a brief current injection shifted the phase of the rhythm (Fig.2A). We defined the phase as the normalized time since the first spike in the most recent volley of spikes. Reflecting the strictly positive PRC of the individual integrate-fire neurons, without heterogeneity (*CV* = 0) external inhibition always delayed the rhythm, independent of the timing of the pulse. In contrast, in heterogeneous networks the rhythm could be advanced if the same inhibitory perturbation was applied shortly after the first spikes in the spike volley (*ϕ*_*inh*_ *>* 0). Increasing the neuronal heterogeneity enhanced this phase advance, since it shifted the within-network inhibition driven by the leading neurons to earlier times, while it delayed the lagging neurons. As a result, for the same external perturbation, a larger fraction of neurons that would spike in the absence of the external inhibition was sufficiently delayed to have their spikes be suppressed by the within-network inhibition (cf. Fig.1B), reducing the within-network inhibition and with it the time to the next spike volley. To keep the frequency of the unperturbed network fixed in Fig.2A, we reduced the tonic input with increasing heterogeneity, which enhanced the phase advance. However, even if the tonic input was kept fixed, the phase advance increased with heterogeneity (Suppl. Fig.S2).

**Figure 2:**
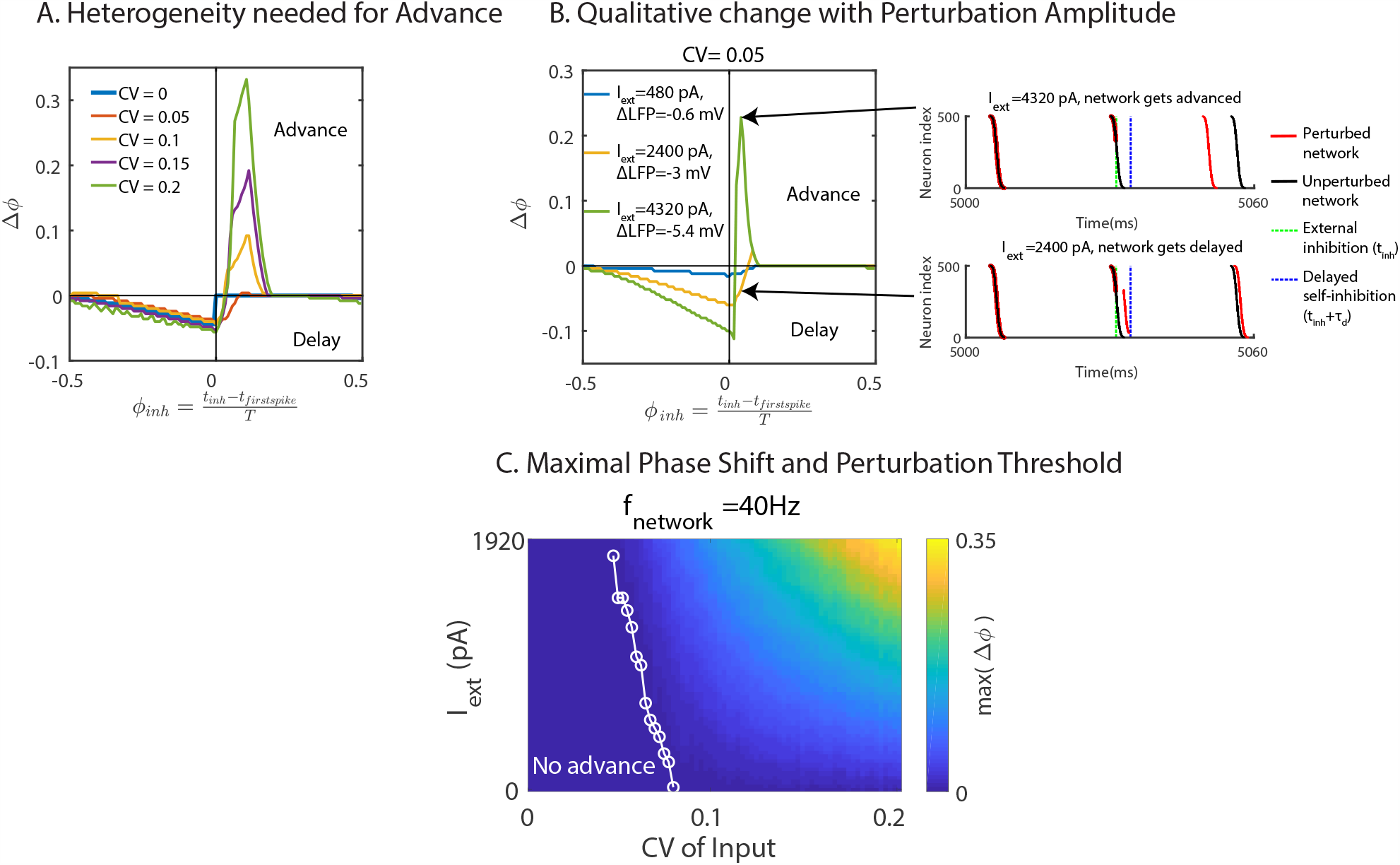
fmPRC of heterogeneous ING network. (A) Phase shift in response to inhibition for different neuronal heterogeneity but fixed natural frequency (*f*_*network*_ = 40Hz). The paradoxical phase advance increased with neuronal heterogeneity. (B) fmPRC changed qualitatively with the amplitude of the perturbation. Left: fmPRC for three different perturbation amplitudes. Right: raster plot of spikes without (black) and with (red) external inhibition. Top: strong perturbation advanced the network. Bottom: weak perturbation applied at the same time as in the top figure. The network was delayed. (C) Maximal phase advance as a function of neuronal heterogeneity and external inhibition strength. The threshold of the inhibition amplitude to obtain an advance decreased with heterogeneity (white line). *f*_*network*_ was kept constant (*f*_*network*_ = 40Hz). In (A)-(C), perturbations were made with a square-wave inhibitory current pulse with duration 0.1 ms to each interneuron. In (A), the amplitude of the current was 1600 pA, resulting in a 2 mV rapid hyperpolarization.

For weak heterogeneity the paradoxical phase response occurred only for sufficiently strong perturbations, i.e. it did not arise in the infinitesimal macroscopic PRC (imPRC). Thus, the phase response changed qualitatively as the amplitude of the perturbation was strong enough to delay the spikes of sufficiently many slow neurons until the self-inhibition of the network set in and suppressed their spikes (Fig.2B). As the CV of the neurons was increased, the dispersion was large enough that the spikes of the lagging neurons were suppressed by the self-inhibition of the network even in the absence of an external perturbation. Above that threshold value of CV the paradoxical phase response occurred even for infinitesimal perturbations (Fig.2C).

The paradoxical phase response was robust with respect to changes in the natural frequency of the network, the coupling strength, and the effective synaptic delay, as long as the rhythm persisted. The paradoxical phase advance increased with decreasing natural frequency of the network, since the inhibition had a stronger effect for lower mean input strength (Fig.3A). Changing the within-network coupling strength by a factor of 2 up or down did not substantially affect the paradoxical phase response (Fig.3B) nor the strength of the rhythm (Fig.3C). Even without explicit synaptic delay (*τ*_*d*_ = 0), the effective delay given by the double-exponential synaptic interaction was sufficient to render a paradoxical response (Fig.3D). However, when this effective delay was reduced by decreasing the rise time 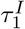 of the synaptic current, the rhythm itself developed a strong subharmonic component and eventually disintegrated (Fig.3E). In the subharmonic regime the paradoxical phase advance alternated in consecutive cycles of the rhythm.

**Figure 3:**
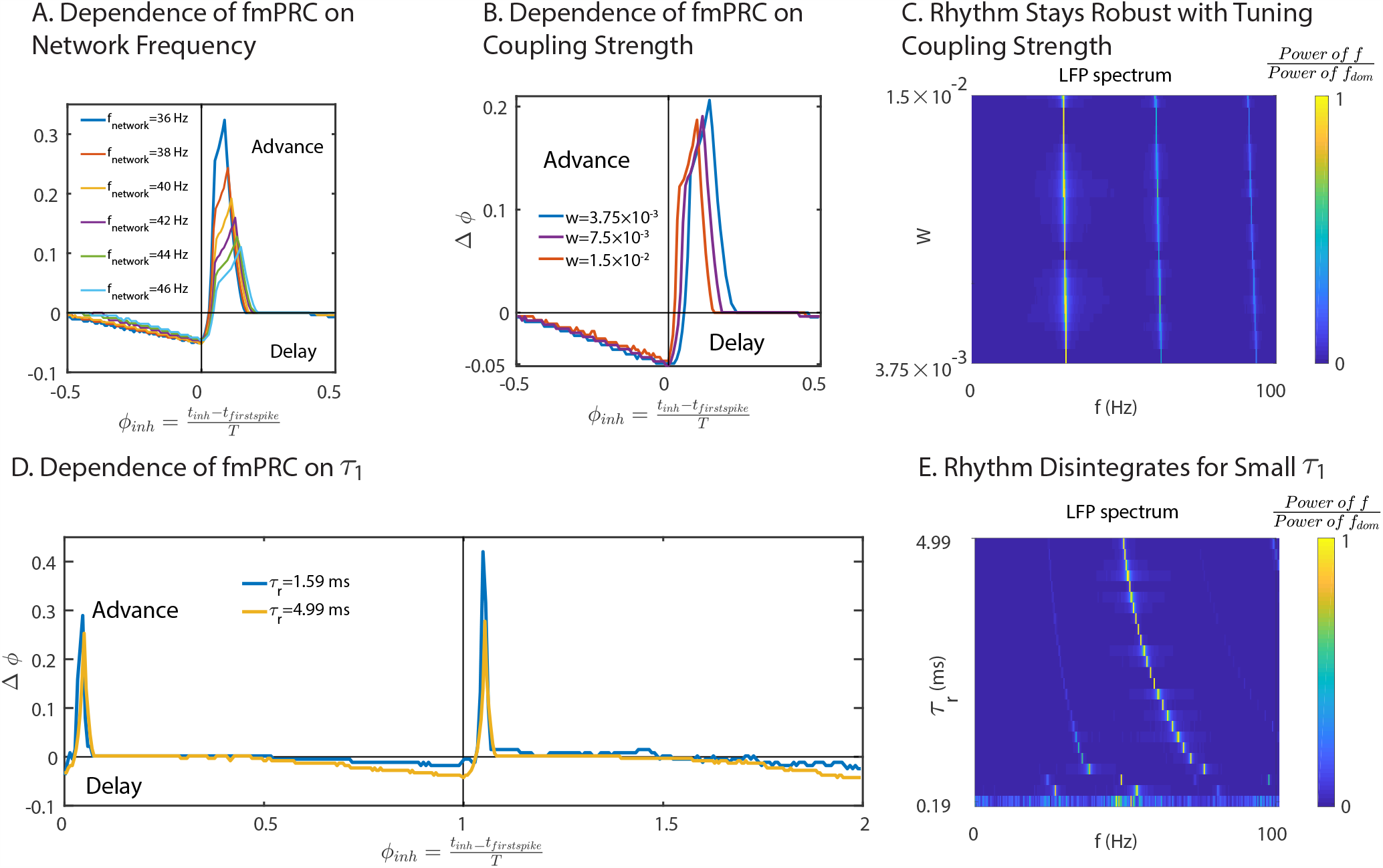
The paradoxical phase response of a heterogeneous ING network is robust. (A) The phase advance of the fmPRC decreased with the natural frequency (*CV* ^(*I*)^ = 0.15). (B) The fmPRC did not depend sensitively on the within-network coupling strength *W* (*CV* ^(*I*)^ = 0.15, *I*^(*I*)^ =15.8 pA). (C) The Fourier spectrum of the LFP as a function of the coupling strength *W* . Parameters as in (B). (D) Paradoxical phase response in the absence of an explicit delay, *τ*_*d*_ = 0, for different effective synaptic delays due to different synaptic rise times 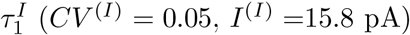. For low 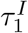 (blue curve), the shift alternated in subsequent cycles reflecting the subhamornic nature of the rhythm. (E) The Fourier spectrum of the LFP as a function of the effective synaptic delay (synaptic time constant of rise 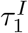). With decreasing 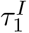, a subharmonic peak emerged and eventually the rhythm disintegrated. Parameters as in D. In (A), (B) and (D), perturbations were made with a square-wave inhibitory current pulse with duration 0.1 ms to each interneuron. In (A) and (B), the amplitude of the current was 1600 pA, resulting in a 2 mV rapid hyperpolarization. In (D), the amplitude of the current was 400 pA, resulting in a 0.5 mV rapid hyperpolarization.

**Figure 4:**
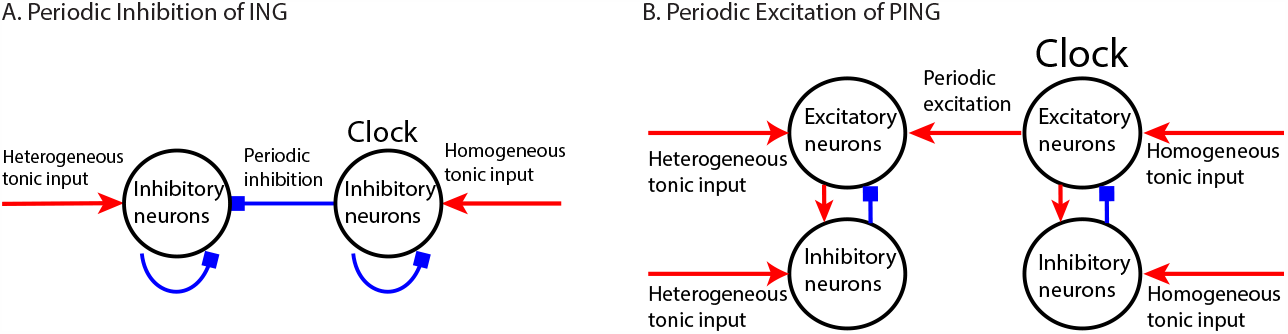
Sketch of computational models. (A) ING rhythm receives periodic inhibitory input generated from another ‘clock’ ING rhythm. (B) PING rhythm receives periodic excitatory input by its E-population generated from another ‘clock’ PING rhythm.

In [13, 27] the exact reduction of all-to-all coupled heterogeneous networks of quadratic integrate-fire neurons to 2 coupled ordinary differential equations for each network [23] has been used to obtain the infinitesimal macroscopic phase-response curve (imPRC) for ING and PING networks. They obtained biphasic response only if the excitatory perturbation was applied to the population of inhibitory neurons; for perturbations to the excitatory neurons they found only monophasic response (type-I). This is presumably due to the lack of a delay in the single-exponential synaptic interactions used in [13, 27].

### Enhancing entrainment of ING-rhythms through network heterogeneity

In order to allow communication by coherence [11, 32], the rhythms in different brain areas need to be sufficiently phase-locked with each other. As a simplification of two interacting *γ*-rhythms, we therefore investigated the ability of the rhythm in a network to be entrained by a periodic external input, particularly focusing on the possibly facilitating role of neuronal heterogeneity. Motivated by the paradoxical phase response induced by the heterogeneity, we addressed, in particular, the question whether an ING network can be sped up by inhibition to entrain it with a faster network.

The network considered here was the same as that used to analyze the fmPRC. The within-network interaction was an all-to-all inhibition with synaptic delay *τ*_*d*_, resulting in a rhythm with natural frequency *f*_*natural*_, Each neuron received heterogeneous input *I*_*bias*_ and inhibitory periodic pulses with frequency *f*_*clock*_ . The latter can be considered as the output of another ING-network and were, in fact, generated that way (Fig.4). We refer to this external input as the ‘clock’. The detuning Δ*f* = *f*_*clock*_ − *f*_*natural*_ was a key control parameter.

For periodic input the fmPRC allows the definition of an iterated map describing the response of the network. For periodic *δ*-pulses that map is shown in Fig.5A. For positive detuning, i.e. when the clock is faster than the network, entrainment requires that the phase response is paradoxical in order for the rhythm to be sped up by the inhibition. If the heterogeneity and the resulting phase response are sufficiently large, the maximum of the iterated map crosses the diagonal, generating a stable and an unstable fixed point. The former is the desired entrained state.

As the detuning is increased the iterated map is shifted downward. This can decrease the slope of the iterated map at the fixed point below −1, destabilizing the fixed point in a period-doubling bifurcation. For periodic pulses comprised of double-exponential inhibitory currents (cf. (2,3)) a rich bifurcation scenario emerged (Fig.5B). Note that the full map is not continuous and not unimodal (cf. first bottom panel of Fig.5B). Nevertheless, for Δ*f <* 7.17 Hz the attractor remains near the unstable fixed point and displays a period-doubling cascade to chaos and multiple periodic windows. For Δ*f >* 7.28 Hz, however, the attractor includes points on both sides of the discontinuity (cf. third bottom panel in Fig.5B).

**Figure 5:**
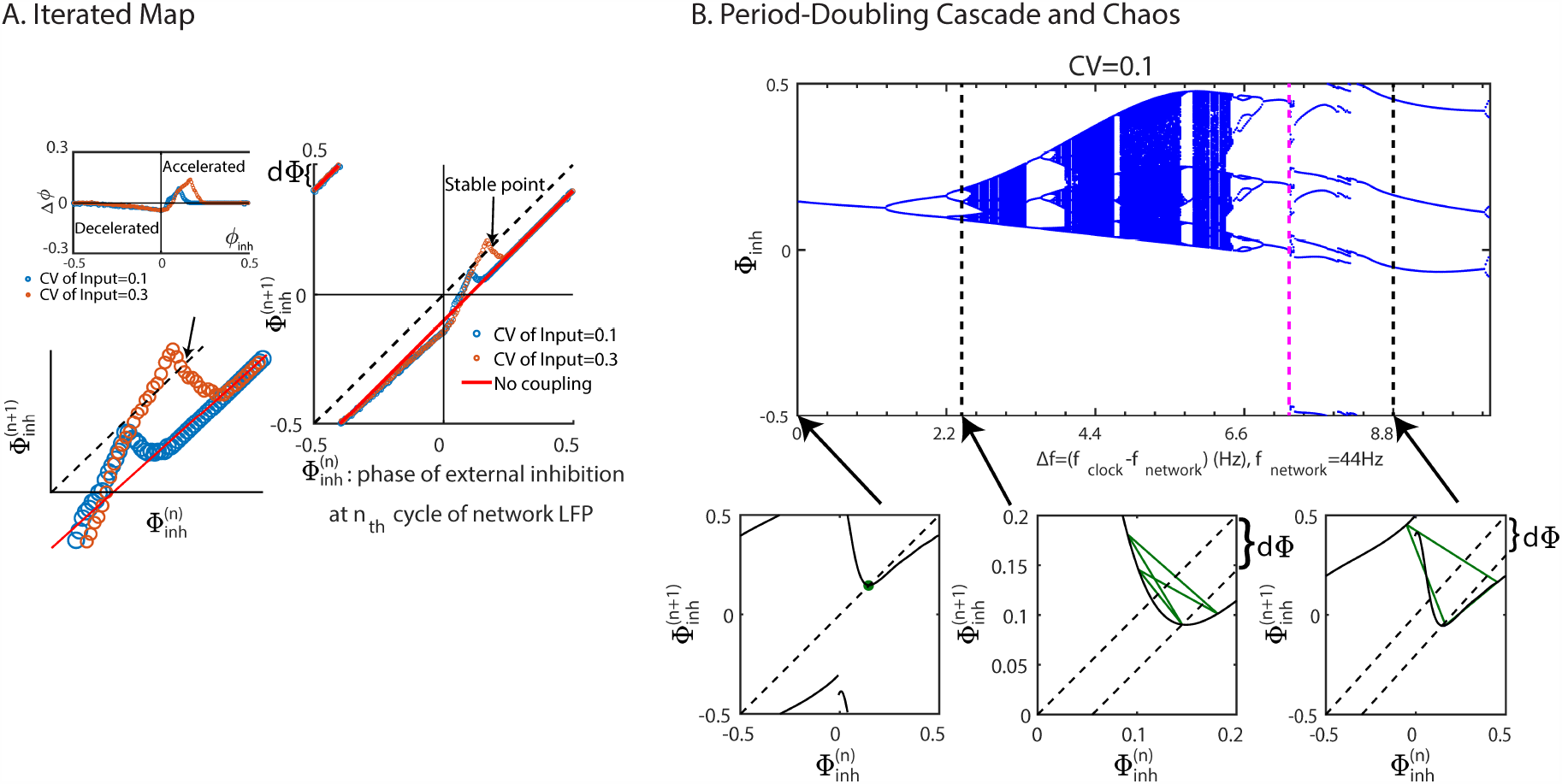
Connection between fmPRC and the synchronization of *γ*-rhythms. (A) Iterated map of Φ_*inh*_. The network can be synchronized by faster periodic inhibiton under sufficiently large advancing phase response. (B) Top: The bifurcation diagram of the iterated map of Φ_*inh*_ with varying detuning Δ*f* . To the right of the magenta dashed line (Δ*f* = 7.28 Hz) the attractors involve points on both sides of the discountinuity of the map. Bottom from left to right: iterated maps of Φ_*inh*_ for Δ*f* = 0, 2.44, 8.8 Hz. The distance between the diagonal and subdiagonal line represents the detuning between the network and periodic input. In (A), the fmPRC was determined for a *δ*-pulse perturbation, in (B) for a double-exponential inhibitory current (cf. (2,3)) was used as in Fig.6.

Having clarified the role of the fmPRC in the network’s synchronizability and ability to phase-lock, we investigated the role of neuronal heterogeneity in more detail (Fig.6). To do that, we adjusted for each value of the input heterogeneity the mean input strength *I*^(*I*)^ so as to keep the natural frequency *f*_*network*_ constant (*f*_*network*_ = 44 Hz). Then we determined the extent of synchronization and phase-locking of the network under the influence of periodic inhibitory input as a function of the detuning Δ*f* and network heterogeneity *CV* . As shown above, the fmPRC of a heterogeneous network could be biphasic with the amplitude of the paradoxical phase response increasing with neuronal heterogeneity. Expecting that for sufficiently large heterogeneity an ING-rhythm could be accelerated by a faster periodic inhibition, we tested phase-locking predominantly for positive detuning, corresponding to *f*_*clock*_ *> f*_*network*_. We first investigated how neuronal heterogeneity affected the synchronization by comparing the dominant frequency *f*_*dom*_ in the Fourier spectrum of the network’s LFP with *f*_*clock*_. In Fig.6A, the color hue indicates the ratio *f*_*dom*_ : *f*_*clock*_. For small heterogeneity, *f*_*dom*_ was a rational multiple of *f*_*clock*_ that depended on the detuning, while for sufficiently large CV the network became synchronized in the sense that *f*_*dom*_ = *f*_*clock*_ (yellow). The range of Δ*f* allowing synchronization became wider with increasing neuronal heterogeneity, implying that the neuronal heterogeneity enhanced the synchronization of the ING-rhythm. However, note that *f*_*dom*_ = *f*_*clock*_ did not imply a perfectly synchronized or a 1:1 phase-locked state. In fact, various different subharmonic responses arose: example 2 shows a period-4 state, while in example 3 the dynamics were actually chaotic (Fig.6A) even though *f*_*dom*_ = *f*_*clock*_. Motivated by these observations, we divided the states into three types:

**Figure 6:**
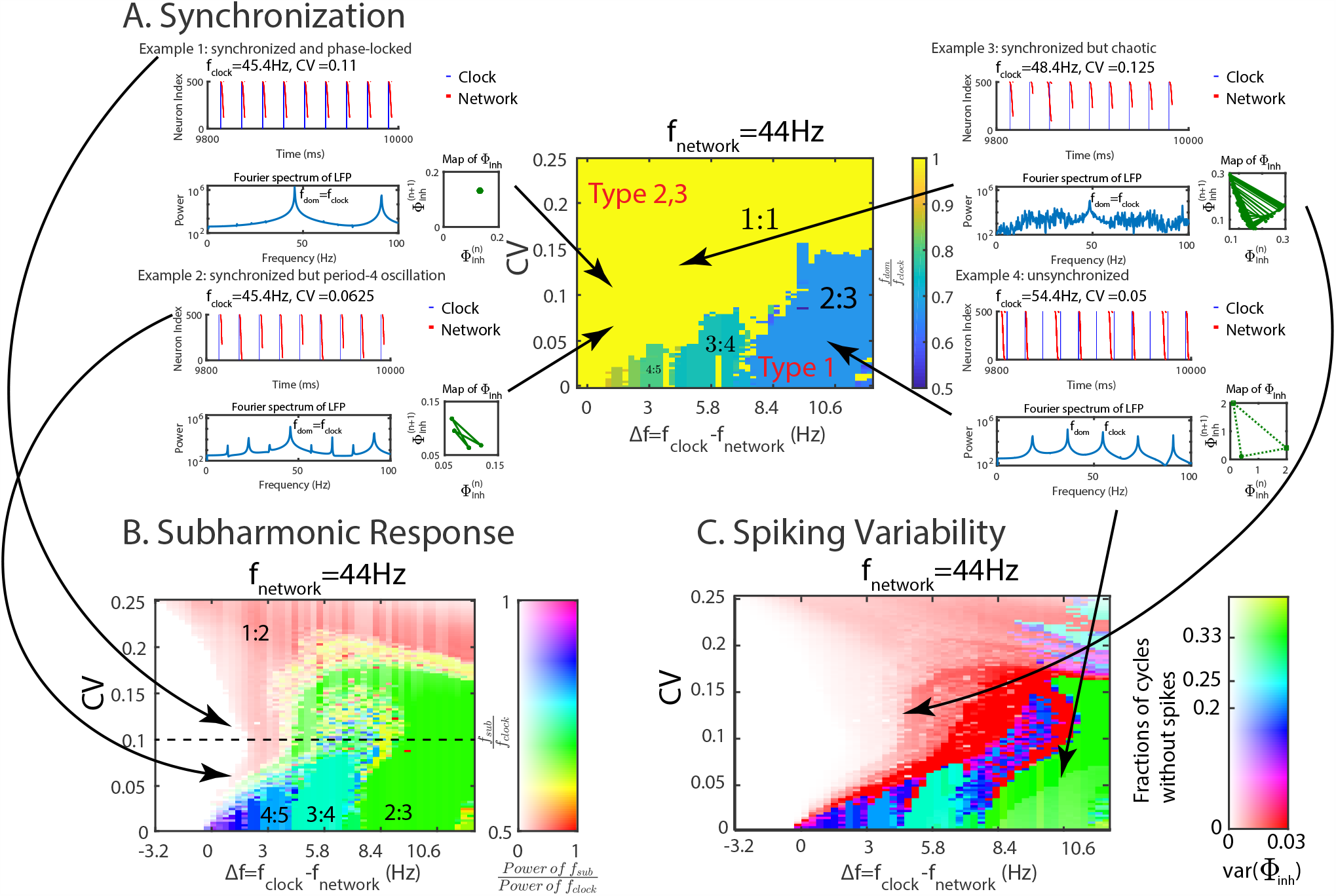
Network heterogeneity enhances synchronization and phase-locking of periodically driven ING rhythm. (A) Synchronization quantified using *f*_*dom*_ : *f*_*clock*_ with *f*_*dom*_ and *f*_*clock*_ being the dominant frequencies of the Fourier spectrum of the LFP of the network and the clock, respectively. The neuronal heterogeneity enhanced the synchronization by shifting *f*_*dom*_ to *f*_*clock*_. Example 1: Synchronized with 1:1 phase-locking. Example 2: Synchronized with subharmonic response (period 4). Example 3: synchronized with subharmonic response (chaotic). Example 4: Not synchronized. Squares and dashed lines in the map of Φ_*inh*_ indicate clock cycles in which the network did not spike (Φ_*inh*_ was arbitrarily set to 2). (B) Subharmonic response. Color hue and saturation indicate the frequency ratio *f*_*sub*_ : *f*_*clock*_ and the ratio of the Fourier power at these two frequencies. *f*_*sub*_ is the frequency of the dominant peak of the network power spectrum that satisfies *f*_*sub*_ *< f*_*clock*_. The power ratio is capped at 1. Dashed line marks the value of input heterogeneity used in Fig.5B. (C) Spiking variability and *var*(Φ_*inh*_) as a function of neuronal heterogeneity and detuning. Color hue indicates the fraction of clock cycles without spikes in the network. In particular, red indicates that the network spikes in every cycle. Color saturation indicates *var*(Φ_*inh*_). The neuronal heterogeneity enhances the tightness of the phase-locking.

- Type 1: *f*_*dom*_ ≠ *f*_*clock*_, not synchronized, not phase-locked (Fig.6 example 4).
- Type 2: *f*_*dom*_ = *f*_*clock*_ with subharmonic response, might be poorly phase-locked (Fig.6 example 3) or displaying rational ratio phase-locking (Fig.6 example 2).
- Type 3: *f*_*dom*_ = *f*_*clock*_, no subharmonic response, 1-to-1 phase-locking (Fig.6 example 1).

The phase diagram Fig.6A does not differentiate between types 2 and 3. It only shows that neuronal heterogeneity enhanced the synchronization of the network by shifting *f*_*dom*_ to *f*_*clock*_. Therefore, we studied whether neuronal heterogeneity also enhanced the synchronization by weakening the subharmonic response and changing the synchronized state from type 2 to type 3, as well as whether the dynamics of the fmPRC shown in the bifurcation diagram Fig.5B could predict the phase relationship between the network and the clock. Using the same simulation setup as in Fig.6A, the subharmonic response is shown in Fig.6B. The color hue indicates the ratio *f*_*sub*_ : *f*_*clock*_, where *f*_*sub*_ is the frequency of the dominant peak of the LFP power spectrum that satisfies *f*_*sub*_ *< f*_*clock*_. The color saturation gives the ratio of the powers at *f*_*sub*_ and *f*_*clock*_ (capped at 1). Thus, over most of the range of positive detuning and neuronal heterogeneity tested, the fading-away of the color with increasing heterogeneity reveals that the neuronal heterogeneity weakened the subharmonic response. Over a small range of positive detuning, increasing neuronal heterogeneity from small values induced perfect synchronization (type 3) by weakening the subharmonic response with frequency ratio *f*_*sub*_ : *f*_*clock*_ = 1 : 2; the system traversed a continuous period-doubling bifurcation in reverse, with type 2 (red) giving way to type 3 (white). Together with Fig.6A, this showed that neuronal heterogeneity could enhance the synchronization both by making *f*_*dom*_ = *f*_*clock*_ (from type 1 to type 2) and by weakening the subharmonic response (from type 2 to type 3). The range of detuning where increasing heterogeneity induced a type 3 synchronization became wider for larger synaptic delay within the network (Suppl. Figure S3). Note that the bifurcation diagram (Fig.5B) based on the fmPRC agrees well with the subharmonic response marked along the dashed line at *CV* = 0.1 in Fig.6B, suggesting that the fmPRC can well predict the subharmonic response and persistent phase response of the network.

In addition to enhancing the frequency synchonization, neuronal heterogeneity was also able to increase the tightness of the phase-locking. Over most of the parameter regime investigated, the variance of the phase of the network relative to the periodic input (*var*(Φ_*inh*_)) decreased with increasing heterogeneity, as indicated by the decrease in the color saturation in Fig.6C. In fact, for detuning between 0 Hz and 2 Hz the heterogeneity reduced *var*(Φ_*inh*_) to 0 (white), corresponding to the 1:1 phase-locked state. Even for the 1:2 phase-locked state (cf. the red area in Fig.6B) *var*(Φ_*inh*_) was very small for a range of heterogeneity and detuning (2 Hz to 4 Hz), indicating tight phase locking. Except for type-3 synchronized states the size of the spike volleys varied between clock cycles. In fact, over wide ranges of the parameters the network did not spike in each of the clock cycles, as indicated by the color hue in Fig.6C, which gives the fraction of cycles with no network spikes (e.g. Fig.6 example 4).

### Paradoxical phase response and entrainment of PING rhythms

Many *γ*-rhythms involve not only inhibitory neurons, but arise from the mutual interaction of excitatory (E) and inhibitory (I) neurons (PING rhythm) [33]. The key elements to obtain a paradoxical phase response and the ensuing enhanced synchronization are self-inhibition within the network, neuronal heterogeneity and effective synaptic delay. Since in PING rhythms the connections from E-cells to I-cells and back to the E-cells form an effective self-inhibiting loop, we asked whether PING-rhythms can exhibit behavior similar to the behavior we identified for ING-rhythms.

Considering a PING-rhythm generated by an E-I network comprised of integrate-fire neurons, we first studied its fmPRC. To avoid that all I-cells receive identical input and therefore spike as a single unit, the I-cells received, in addition to the excitation from the E-cells, heterogeneous, tonic, Gaussian-distributed subthreshold input with mean *I*^(*I*)^ = 36 pA and *CV* ^(*I*)^= 0.167. The phase response of the network was probed by applying an identical external excitatory perturbation to all E-cells and recording the resulting phase shift of the LFP (cf. eqs.(7,8)) of the E-population, averaged across 500 realizations of the subthreshold input to the I-cells (Fig.7A). More specifically, the perturbations consisted of a square-wave excitatory current pulse with amplitude 76 pA and duration 0.1 ms to each E-cell, resulting in a 2 mV rapid depolarization. Without neuronal heterogeneity the external excitation always advanced the phase of the rhythm resulting in an fmPRC that was strictly positive. In the heterogeneous case, however, the PING rhythm exhibited a paradoxical phase response, whereby the collective rhythm was delayed while the individual neurons were advanced by the excitation. The delay was caused by the increase of self-inhibition within the network that was generated by the additional spikes in the E-population, which in turn drove additional spikes in the I-population. In contrast to the fmPRC of the ING-rhythm, this paradoxical phase response was not monotonic in the heterogeneity. While weak heterogeneity resulted in strong delay, the delay decreased with increasing intermediate CV-values and only increased again for larger CV (Fig.7B left top). This non-monotonicity arose because we kept the frequency of the network constant as we increased its heterogeneity. This required a decrease in the tonic input to the E-cells with increasing heterogeneity. For the stronger tonic input used for weak heterogeneity the same external perturbation elicited more additional spikes than it did for strong heterogeneity where the tonic input was weaker (cf. titles of subpanels of Fig.7B). The total number of spikes occurring in each cycle of the unperturbed network also decreased with increasing heterogeneity. Consequently, the relative change in the number of spikes induced by the perturbation was non-monotonic in the heterogeneity. As a result, the relative change in the inhibitory synaptic conductance resulting from the perturbation and with it the phase delay was also non-monotonic.

**Figure 7:**
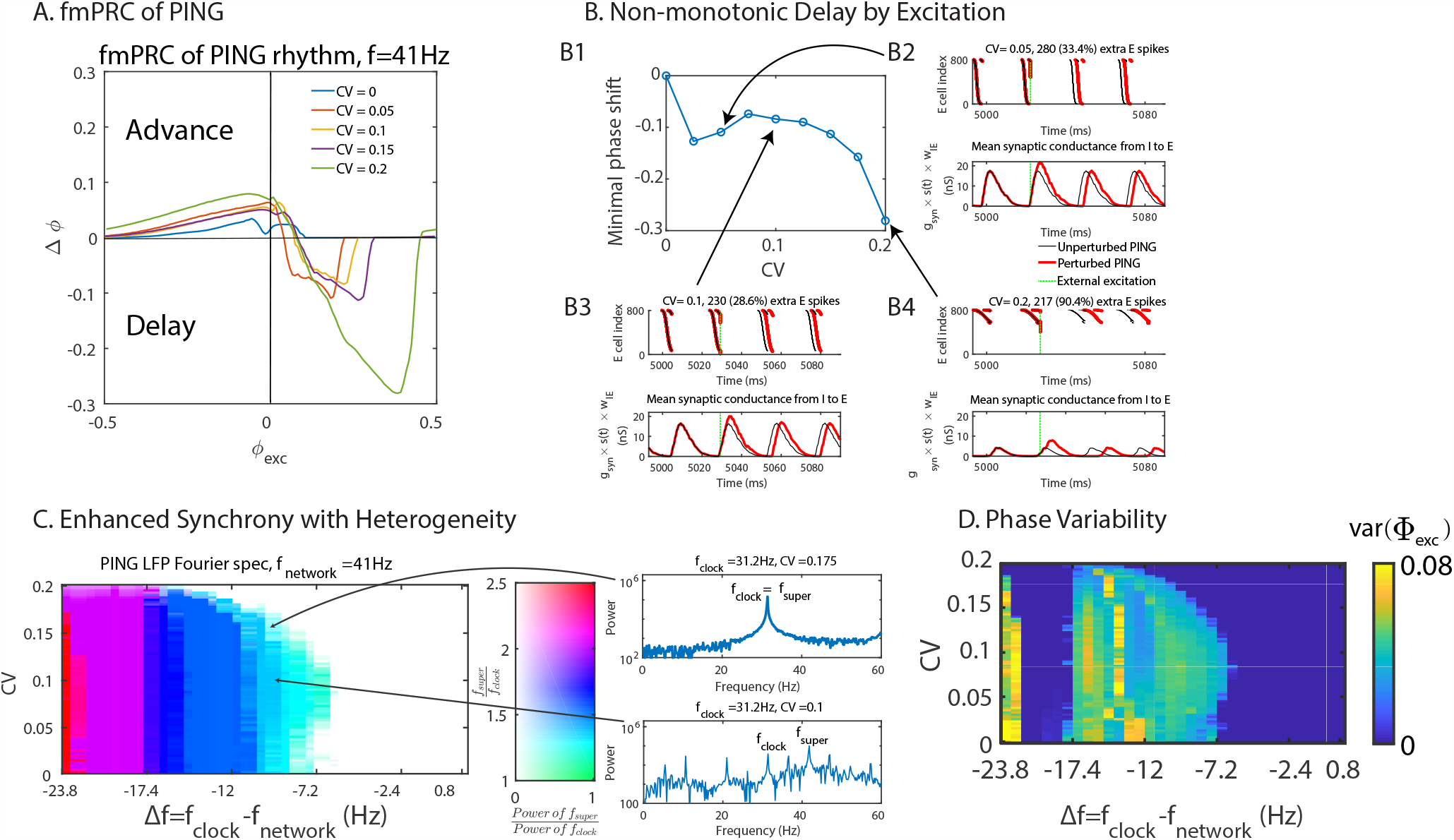
Network heterogeneity enhances the synchronization and the tightness of phase-locking of the PING rhythm. (A) fmPRC of PING networks with constant natural frequency (*f*_*network*_ = 41 Hz) but different neuronal heterogeneity. Only with neuronal heterogeneity the phase was delayed by the excitation. (B) Non-monotonicity of the paradoxical delay with constant natural frequency (*f*_*network*_ = 41 Hz). B2-4: Top: raster plot of spikes in E-population (input strength increased with cell index). Bottom: mean inhibitory synaptic conductance within the PING network. The titles show the absolute and relative increase in spike number (B2: *CV* = 0.05, B3: *CV* = 0.1, B4: *CV* = 0.2). (C) Subharmonic response of the PING rhythm with periodic excitation as function of neuronal heterogeneity and detuning. *f*_*network*_ was fixed at 41 Hz. Color hue and saturation indicate the frequency ratio and power ratio at the frequencies *f*_*super*_ and *f*_*clock*_ of the E-population’s LFP. *f*_*super*_ was the frequency of the dominant peak of the LFP power spectrum that satisfies *f*_*super*_ *> f*_*clock*_. The power ratio was capped at 1. Generally, the neuronal heterogeneity enhanced the synchronization of the PING rhythm by weakening subharmonic response. (D) The tightness of the phase-locking (*var*(Φ_*exc*_)) as a function of neuronal heterogeneity and detuning. The neuronal heterogeneity enhanced the tightness of the phase-locking. For Δ*f* ∈ [−22Hz, −17.4Hz] the clock was twice as fast as the network, resulting in vanishing *var*(Φ_*exc*_).

As for the ING rhythm, we investigated the role of neuronal heterogeneity in the synchonizability and the ability of phase-locking of coupled PING rhythms. In analogy to the ING-case, we considered the case of the E-population of a PING network receiving periodic excitation generated by a clock PING network (Fig.4B). As before, we adjusted the tonic input strength to the E-population to keep the natural frequency of the network constant as we changed its heterogeneity (*f*_*network*_ = 41Hz). To probe the impact of the paradoxical phase response on the synchronization we focused on negative detuning for which the periodic external excitation needed to slow down the network in order to achieve phase-locking. Indeed, with increasing heterogeneity the network could become synchronized with the slower clock over a larger ranger of the detuning as indicated by the fading saturation of the color in Fig.7C. Here the color hue indicates the ratio *f*_*super*_ : *f*_*clock*_, where *f*_*super*_ was determined as the frequency with the most power among the frequencies higher than *f*_*clock*_ in the Fourier spectrum of the E-population’s LFP. The color saturation indicates the ratio of the power at the frequencies *f*_*super*_ and *f*_*clock*_. Thus, a color hue closer to green (*f*_*super*_ : *f*_*clock*_ = 1 : 1) or with a lower saturation implies better synchronization. By observing how the width of the range of detuning allowing synchronization varied with neuronal heterogeneity, we concluded that generally, the synchronizability of PING rhythm was enhanced by the neuronal heterogeneity by weakening subharmonic response. Note that for *CV* ∈ [0, 0.1] the synchonizability of the PING rhythm decreased slightly with neuronal heterogeneity. This was consistent with the nonmonotonicity exhibited by the fmPRC seen in Fig.7B. The neuronal heterogeneity played a similar role in the tightness of the phase-locking as in the synchronizability (Fig.7D).

## 3 Discussion

In this paper we have analyzed the response of collective oscillations of inhibitory and of excitatory-inhibitory networks of integrate-fire neurons to external perturbations. For ING- and PING-rhythms we have shown that the combination of neuronal heterogeneity and effective synaptic delay can qualitatively change the phase response compared to the phase response of the individual neurons generating the rhythm. Thus, perturbations that delay the I-cells can paradoxically advance the ING-rhythm and perturbations that advance the E-cells can delay the PING-rhythm. As a result, the macroscopic phase-response curve for finite-amplitude perturbations (fmPRC) of the rhythm can change sign as the phase of the perturbation is changed (type-II), even though the PRC of all individual cells is strictly positive (type-I). This change of the fmPRC enhances the ability of the *γ*-rhythm to synchronize with other rhythms.

The key element of the mechanism driving the paradoxical phase response and the enhanced synchronization is the cooperation of the external perturbation and the effectively delayed within-network inhibition. In the ING-network a suitably timed external perturbation delays the lagging − but not the early - neurons sufficiently to allow the within-network inhibition triggered by the early neurons to keep the lagging neurons from spiking. This reduces the overall within-network inhibition and with it the duration of the cycle. Thus, the perturbation modifies the internal dynamics of the rhythm, which leads to changes in the phase of the rhythm that can dominate the immediate phase change the perturbation induces. The situation is somewhat similar to that investigated in [18]. There it had been pointed out that an external perturbation of a collective oscillation can lead to changes in its phase in two stages: i) an immediate change of the phases of all oscillators as a direct result of the perturbation and ii) a subsequent slower change in the collective phase resulting from the convergence of the disturbed phases back to the synchronized state. That analysis was based on a network of phase oscillators and could therefore not include a key element of our results, which is the perturbation-induced change in the number of neurons that actually spike and the resulting change in the within-network inhibition that results in a change of the period of the rhythm. As discussed in [31, 34], for ING-rhythms such a change in the number of spiking neurons underlies also the enhanced phase-locking found in [30].

Going beyond ING-rhythms, we showed that PING-rhythms can also exibit a paradoxical phase response *via* a mechanism that is analogous to that of ING-rhythms. For that analysis we have focused on excitatory-inhibitory networks with only connections between but not within the excitatory and inhibitory populations. For excitatory inputs to the excitatory cells to generate a paradoxical phase response it is necessary that the additional spikes of the excitatory neurons that are caused by the external perturbation induce additional spikes of the inhibitory neurons. This behavior arises if the inibitory population is also allowed to be heterogeneous. Moreover, the within-network inhibition has to be strong enough to be able to suppress the spiking of lagging excitatory neurons. This is, e.g., found in mice piriform cortex, where principal neurons driven by sensory input from the olfactory bulb arriving early during a sniff recruit inhibitory interneurons via long-range recurrent connections, resulting in the global, transient suppression of subsequent cortical activity [35]. A characteristic feature of the paradoxical phase response of the PING rhythm is the extended cycle time following enhanced activation of the excitatory cells. A strong such correlation between the cycle time and the previous LFP amplitude has been observed for the *γ*-rhythm in hippocampus [36]. To assess whether this rhythm exhibits paradoxical phase response would require comparing the macroscopic phase response [37] with that of indvidual participating neurons.

In order for the global perturbation to affect the various neurons differently, they have to be at different phases in their cycle. Our analysis suggests that the specific cause for this heterogeneity in the spike times does not play an important role. Indeed, as shown in [29], even fluctuating heterogeneities that are generated by noise rather than static heterogeneities reflecting intrinsic properties of neurons can enhance the synchronization of multiple *γ*-rhythms in interconnected networks of identical neurons. Note that the noise driving this synchronization is uncorrelated across neurons. The analysis of that system revealed the same mechanism at work as the one identified here.

In various previous analytical and computational analyses it has been found that the dynamics of the macroscopic phase of a collective oscillation can qualitatively differ from that of the microscopic phase. Thus, for interacting groups of noisy identical phase oscillators the macroscopic phases of the groups can tend to lign up with each other, even if all pair-wise interactions between individual oscillators prefer the antiphase state, and vice versa [20]. An analogous result has been obtained for heterogeneous populations of noiseless oscillators [17].

Qualitative changes have also been found in the macroscopic phase response of rhythms in noisy homogeneous networks when the noise level was changed [21, 22, 28]. Using a Fokker-Planck approach for globally coupled excitable neurons, a type-I mPRC was obtained for weak noise, where the rhythm emerges through a SNIC bifurcation, while a type-II mPRC arose for strong noise that led to a Hopf bifurcation [21]. A similar approach was used to obtain the mPRC via the adjoint method for an extension of the theta-model that implements conductance-based synaptic interactions. Again, although individual theta-neurons have a type-I PRC, a type-II mPRC was obtained for the rhythm, which arose in a Hopf bifurcation [22]. This was also the case in an extension to networks of excitable and inhibitory neurons [28].

Thus, results reminiscent of those presented here have been obtained previously. However, the mechanism underlying them was not addressed in detail and remained poorly understood. We expect that our analysis will provide insight into those systems. The key element of the mechanism discussed here is that due to the dispersion of the spike times, which either results from neuronal heterogeneity or noise, the external perturbation enables the within-network inhibition to suppress the spiking of a larger number of neurons than without it. In our system this was facilitated by the delay with which spikes triggered the within-network inhibition, which allowed some neurons to escape its impact in the absence of the external perturbation, but not in its presence. Our analysis showed, however, that the explicit delay is not necessary; the effective delay resulting from a double-exponential synaptic interaction is sufficient. In fact, when reducing that effective delay the paradoxical phase response did not disappear until the delay was so short that the rhythm itself developed a strong subharmonic component and disintegrated.

In this paper we have focused on a specific, very simple neuronal model, the leaky integrate-fire model with conductance-based pulsatile coupling. In previous work on the enhanced synchronization among *γ*-rhythms *via* noise-induced spiking heterogeneity it was demonstrated that this result does not depend sensitively on the neuron type. Comparable results were obtained also with Morris-Lecar neurons for parameters in which the periodic spiking arises from a SNIC-bifurcation, resulting in a type-I PRC as is the case for integrate-fire neurons, but also for parameters for which the spiking is due to a Hopf bifurcation, resulting in a type-II PRC [29]. For networks comprised of heterogeneous neurons with type-II PRC the fmPRC of the collective oscillation is likely to be more complex, since the heterogeneity allows the same input to induce phase shifts with opposite signs for different neurons. However, we expect that the interplay between the within-network inhibition and the external perturbation can again substantially and qualitatively modify the fmPRC by changing the number of neurons participating in the rhythm.

In [29] the results were also found to be robust with respect to significant changes in the network connectivity (random instead of all-to-all) as well as the reversal potential of the inhibitory synapses, as long as the rhythm itself persisted robustly (cf. [38]). In fact, the coupling did not even have to be synaptic; collective oscillations of relaxation-type chemical oscillators that were coupled diffusively were also shown to exhibit noise-induced synchronization. These results suggest that the paradoxical phase response found here arises in a much wider class of macroscopic collective oscillations.

The strong paradoxical phase response that we demonstrated for heterogeneous networks allows their rhythm to synchronize with a periodic external input over a range of detuning that increases sub-stantially with the neuronal heterogeneity. This is reminiscent of computational results for anterior cingulate cortex that investigated networks of excitatory neurons coupled via a common population of inhibitory neurons. There heterogeneity was also found to enhance the synchrony of rhythms, as measured in terms of coincident spikes within 10ms bins [39].

The heterogeneity-enhanced synchrony we have identified suggests that the coherence of *γ*-rhythms emerging in different interacting networks could also be enhanced by neuronal heterogeneity. It has been proposed that the coherence of different *γ*-rhythms, which has been observed to be modified by attention [8], plays an important role in the communication between the corresponding networks [11,32]. Computational studies have shown that the direction of information transfer between networks depends on the relative phase of their rhythms [12, 13], which can be changed by switching between different base states [40, 41]. Whether the enhanced synchrony resulting from neuronal heterogeneity enhances this information transfer is still an open question.

Disrupted *γ*-rhythms have been observed in multiple brain regions in neurological diseases, especially Alzheimer’s disease. Optogenetic and sensory periodic stimulation at *γ*-frequencies has been found to entrain the *γ*-rhythm in hippocampus and visual cortex, respectively, and has resulted in a significant reduction in total amyloid level [42]. Similar neuro-protective effects of entrainment by external *γ*-stimulation have also been found for other sensory modalities [14,43]. This suggests that *γ*-stimulation by sensory input might be a feasible therapeutic approach. Our results suggest a potential role of neuronal heterogeneity in this context.

From a functional perspective, it has been shown that the noise-induced synchronization mentioned above can facilitate certain learning processes [44]. Specifically, a read-out neuron was considered that received input from neurons in two networks *via* synapses that exhibited spike-timing dependent plasticity. The two networks were interacting with each other and each of them exhibited a *γ*-rhythm, albeit at different frequencies. For low noise the two rhythms were not synchronized and the read-out neuron received inputs from the two networks at uncorrelated times. These inputs drove the plasticity inconsistently, leading only to a very slow overall evolution of the synaptic weights, if any. However, for stronger noise the two networks were synchronized, providing a more consistent spike timing that lead to substantial changes in the synaptic weights. As a result, the read-out neuron was eventually only driven by the network that had the larger natural frequency in the absence of the coupling between the networks. It is expected that synchrony by neuronal heterogeneity will have a similar impact.

## 4 Methods

### Neuron model

Both E-cells and I-cells were modeled as leaky integrate-and-fire neurons, each characterized by a membrane potential *V*_*i*_(*t*) satisfying

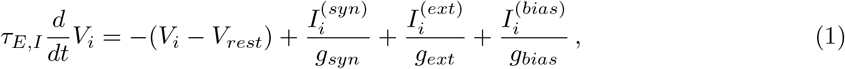

where *V*_*rest*_ is the resting potential and *τ*_*E,I*_ the membrane time constants of the E- and I-cells, respectively. 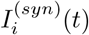 is the total synaptic current that the neuron receives from the other neurons within the network. 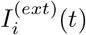 is a time-dependent external input that represents perturbations applied to determine the fmPRC or, in the study of synchronization, the periodic input generated by the clock network. 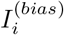 denotes a tonic, neuron-specific excitatory bias current that implements the heterogeneity of the neuron properties. The corresponding conductances are denoted *g*_*syn*_,, *g*_*ext*_, and *g*_*bias*_. Upon the *i*^th^ neuron reaching the spiking threshold *V*_*peak*_, the voltage *V*_*i*_ was reset to the fixed value *V*_*reset*_. Parameters for the neuron were kept fixed throughout all simulations (see Table 1). The local field potential (LFP) of the network was approximated as the mean voltage across all neurons *j* = 1, …*N* in the respective population.

**Table 1:**
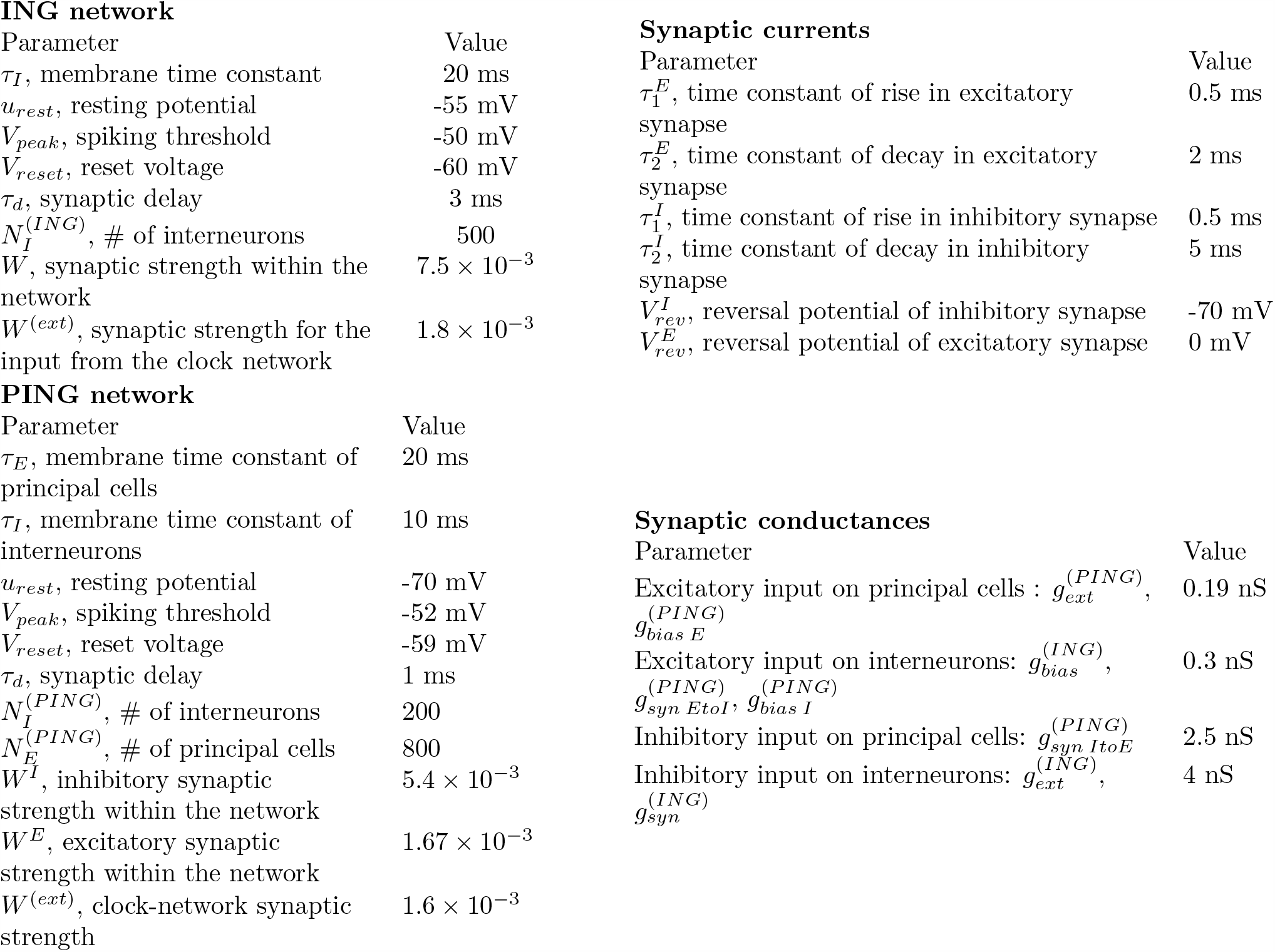
**Parameters used in the model**. Most parameters are based on [29, 45].

ING network
| Parameter | Value |
| --- | --- |
| $\tau_I$, membrane time constant | 20 ms |
| $u_{rest}$, resting potential | -55 mV |
| $V_{peak}$, spiking threshold | -50 mV |
| $V_{reset}$, reset voltage | -60 mV |
| $\tau_d$, synaptic delay | 3 ms |
| $N_I^{(ING)}$, # of interneurons | 500 |
| $W$, synaptic strength within the network | $7.5 \times 10^{-3}$ |
| $W^{(ext)}$, synaptic strength for the input from the clock network | $1.8 \times 10^{-3}$ |

PING network
| Parameter | Value |
| --- | --- |
| $\tau_E$, membrane time constant of principal cells | 20 ms |
| $\tau_I$, membrane time constant of interneurons | 10 ms |
| $u_{rest}$, resting potential | -70 mV |
| $V_{peak}$, spiking threshold | -52 mV |
| $V_{reset}$, reset voltage | -59 mV |
| $\tau_d$, synaptic delay | 1 ms |
| $N_I^{(PING)}$, # of interneurons | 200 |
| $N_E^{(PING)}$, # of principal cells | 800 |
| $W^I$, inhibitory synaptic strength within the network | $5.4 \times 10^{-3}$ |
| $W^E$, excitatory synaptic strength within the network | $1.67 \times 10^{-3}$ |
| $W^{(ext)}$, clock-network synaptic strength | $1.6 \times 10^{-3}$ |

Synaptic currents
| Parameter | Value |
| --- | --- |
| $\tau_1^E$, time constant of rise in excitatory synapse | 0.5 ms |
| $\tau_2^E$, time constant of decay in excitatory synapse | 2 ms |
| $\tau_1^I$, time constant of rise in inhibitory synapse | 0.5 ms |
| $\tau_2^I$, time constant of decay in inhibitory synapse | 5 ms |
| $V_{rev}^I$, reversal potential of inhibitory synapse | -70 mV |
| $V_{rev}^E$, reversal potential of excitatory synapse | 0 mV |

Table 1: **Parameters used in the model.** Most parameters are based on [29,45].
| Parameter | Value |
| --- | --- |
| Excitatory input on principal cells : $g_{ext}^{(PING)}$, $g_{bias E}^{(PING)}$ | 0.19 nS |
| Excitatory input on interneurons: $g_{bias}^{(ING)}$, $g_{syn EtoI}^{(PING)}$, $g_{bias I}^{(PING)}$ | 0.3 nS |
| Inhibitory input on principal cells: $g_{syn ItoE}^{(PING)}$ | 2.5 nS |
| Inhibitory input on interneurons: $g_{ext}^{(ING)}$, $g_{syn}^{(ING)}$ | 4 nS |

### Network model

We studied two types of networks: an ING network and a PING network. The ING network was modeled as an all-to-all inhibitory network of 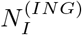 interneurons. The PING network was modeled as a network of 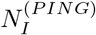 interneurons and 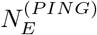 principal cells with all-to-all interneuron-principal and principal-interneuron connections (i.e., without principal-principal and interneuron-interneuron connections). In PING, only principal cells received external input *I*^*ext*^(*t*).

To gain insight into the interaction between two ING rhythms, we considered the simplified situation in which all neurons in the network received strictly periodic input *I*^(*ext*)^, which was generated by another ING network (‘clock’). Similarly, for PING rhythms, the E-cells of the PING network received strictly periodic excitatory input *I*^(*ext*)^ from another PING network through all-to-all connection between their E populations.

### Synaptic currents

We used delayed double-exponential conductance-based currents to model the excitatory and the inhibitory synaptic inputs from neuron *j* to neuron *i*,

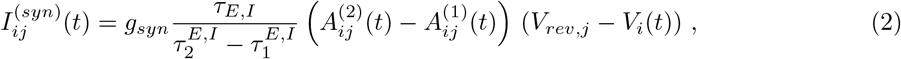

with the two exponentials 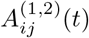 satisfying

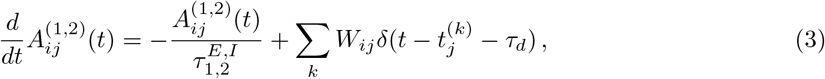

where *V*_*rev,j*_ is the synaptic reversal potential corresponding to the synapse type, *W*_*ij*_ the dimensionless synaptic strength, and *δ* the Dirac *δ*-function. All synapses of the same type (I-I, I-E, E-I) were equally strong. The time constants of 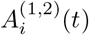 satisfied 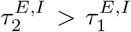 . The synaptic current was normalized to render the time integral independent of the synaptic time constants 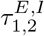. The inhibitory synaptic currents had a slower decay than the excitatory ones (cf. Table 1). We included an explicit synaptic delay *τ*_*d*_ in the model. Every spike of the presynaptic neuron *j* at time 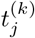 triggered a jump in both 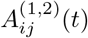, making the synaptic conductance rise continously after a synaptic delay *τ*_*d*_.

External periodic inputs were also modeled as double-exponential conductance-based currents with *g*_*syn*_ in (2,3) replaced by *g*_*ext*_. The time constants and delay were as for the within-network synaptic inputs.

### Heterogeneous tonic input

The bias currents 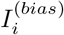 of the ING network were Gaussian distributed around *I*_*mean*_ with a coefficient of variation *CV* and arranged in increasing order, 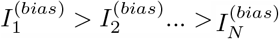. For the PING network, all excitatory neurons received a heterogeneous bias 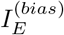 with mean *I*^(*E*)^ and a coefficient of variation *CV* ^(*E*)^. Similarly, the bias currents 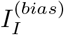 to the inhibitory neurons were characterized by their mean *I*^(*I*)^and their coefficient of variation *CV* ^(*I*)^. Without the excitatory input from principal cells, the voltage of interneurons remained below the spiking threshold. In our investigation of the impact of the neuronal heterogeneity on the phase response and entrainment of the PING rhythm we kept *CV* ^(*I*)^ fixed and varied *CV* ^(*E*)^.

### Macroscopic Phase-response Curve for Finite-Amplitude Perturbations (fmPRC)

#### ING rhythm

For a single ING network, we applied a single inhibitory *δ*-pulse to each neuron 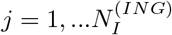 at time *t*_*inh*_ (dashed green line in Fig.1B) and recorded the resulting phase shift Δ*ϕ*. The amplitude of the inhibitory perturbation to each neuron was the same. The phase of the inhibition was defined as

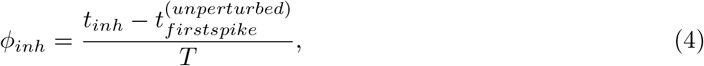

where *T* was the period of the network LFP and 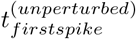 the time of the first spike in the spike volley of the unperturbed network that was closest to *t*_*inh*_. The resulting phase shift Δ*ϕ* was given by

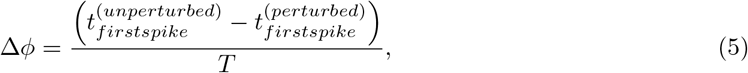

where 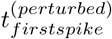 is the time of the first spike in the corresponding volley in the perturbed network. Δ*ϕ* and *ϕ*_*inh*_ were taken to be in the range [−0.5 0.5). Positive Δ*ϕ* indicated that the network was advanced by the perturbation, while negative indicated a delay.

The periodic input (‘clock’) that was used to test the synchronizability of the ING-rhythm was generated by a homogeneous ING network. The phase of the periodic input in the *n*^th^ clock cycle was defined by

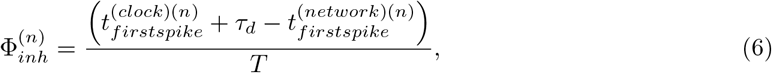

where 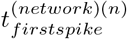 was the time of the first spike in the spike volley of the network in the *n*^th^ cycle and 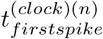 the time of the spike of the clock. In contrast to the definition of *ϕ*_*inh*_ in (4), the definition of 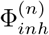 included the delay *τ*_*d*_, since the inhibition generated by the clock arrived with delay *τ*_*d*_ in the network.

#### PING rhythm

To probe the phase response of the PING network we used the same approach as for the ING rhythm, except that we used excitatory instead of inhibitory *δ*-pulses and applied them only to the E-cells. The phase of the excitation *ϕ*_*exc*_ and the resulting phase shift Δ*ϕ* were determined similarly as in the case of the ING rhythm,

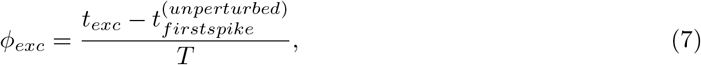

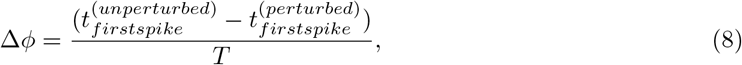

where 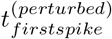 and 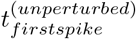 were the times of the first spike in the respective spike volleys of the E-population.

Analogously to 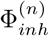, the phase of the periodic input in the *n*^th^ clock cycle was given by

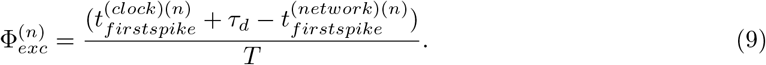

Throughout, the tonic, Gaussian distributed input to the interneurons in the PING network was fixed: *I*^(*I*)^ = 36 pA, *CV* ^(*I*)^ = 0.167.

## Supplementary Information

**Figure S1:**
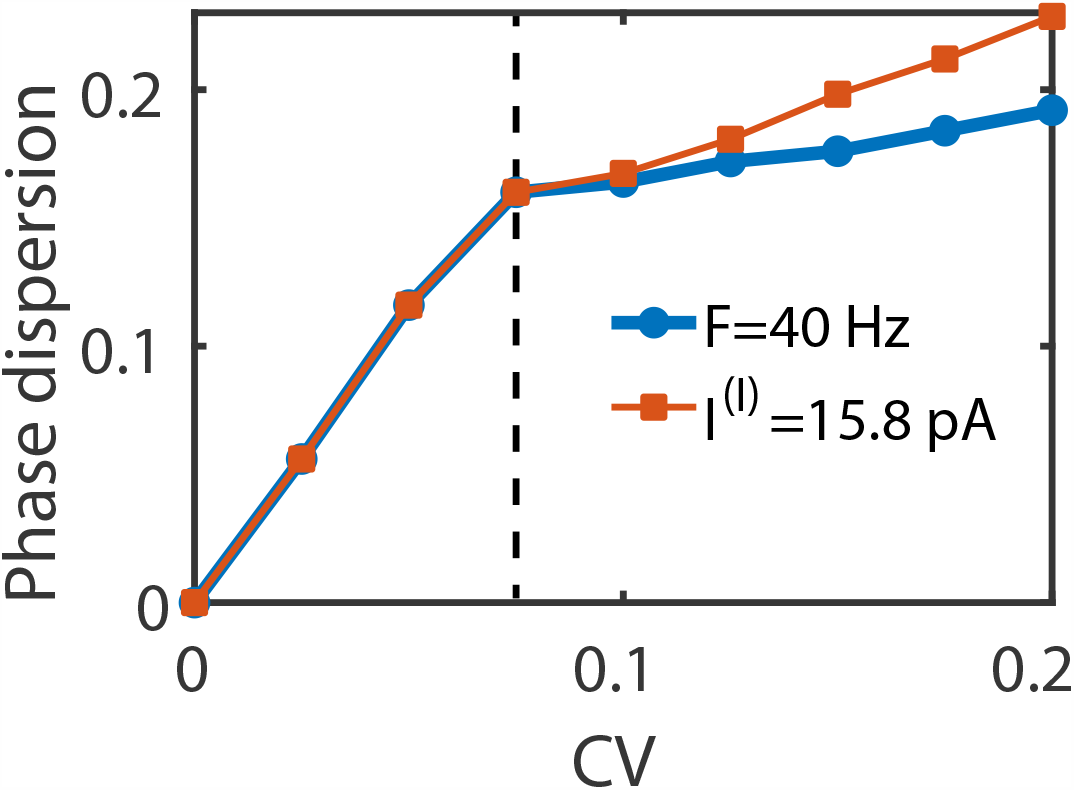
Dependence of the phase dispersion on the heterogeneity of the bias current. The phase dispersion was determined as the time difference between the first and the last spike in the same spike volley normalized by the period. Blue: fixed natural frequency (*f*_*network*_ = 40Hz) for different neuronal heterogeneity. Red: fixed mean input strength (*I*^(*I*)^ =15.8 pA) for different neuronal heterogeneity. For *CV* ≥ 0.075 (dashed line), some neurons spike more than once in a cycle.

**Figure S2:**
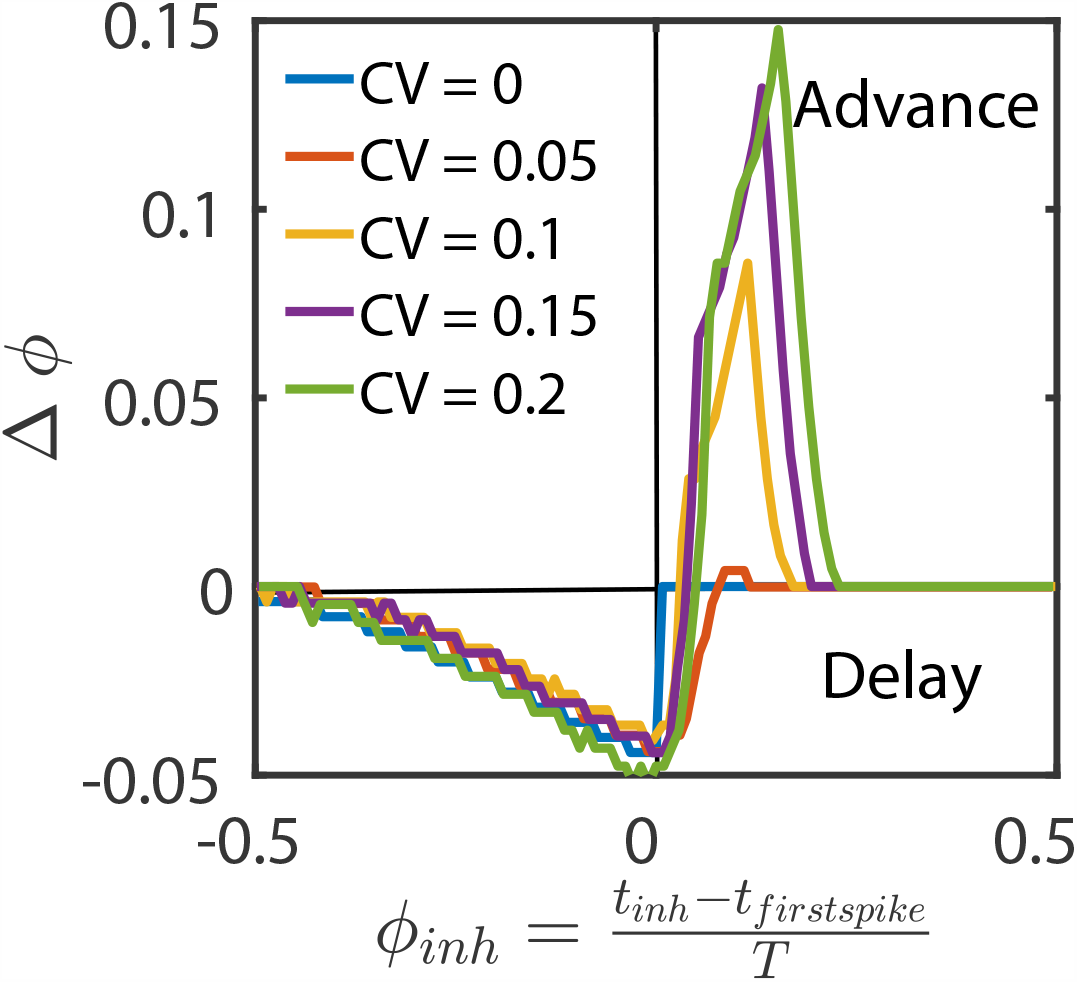
fmPRC of heterogeneous ING networks for fixed steady current (*I*^(*I*)^ =15.8 pA) instead of fixed frequency (cf. Fig.2A). The paradoxical phase advance increased with neuronal heterogeneity.

**Figure S3:**
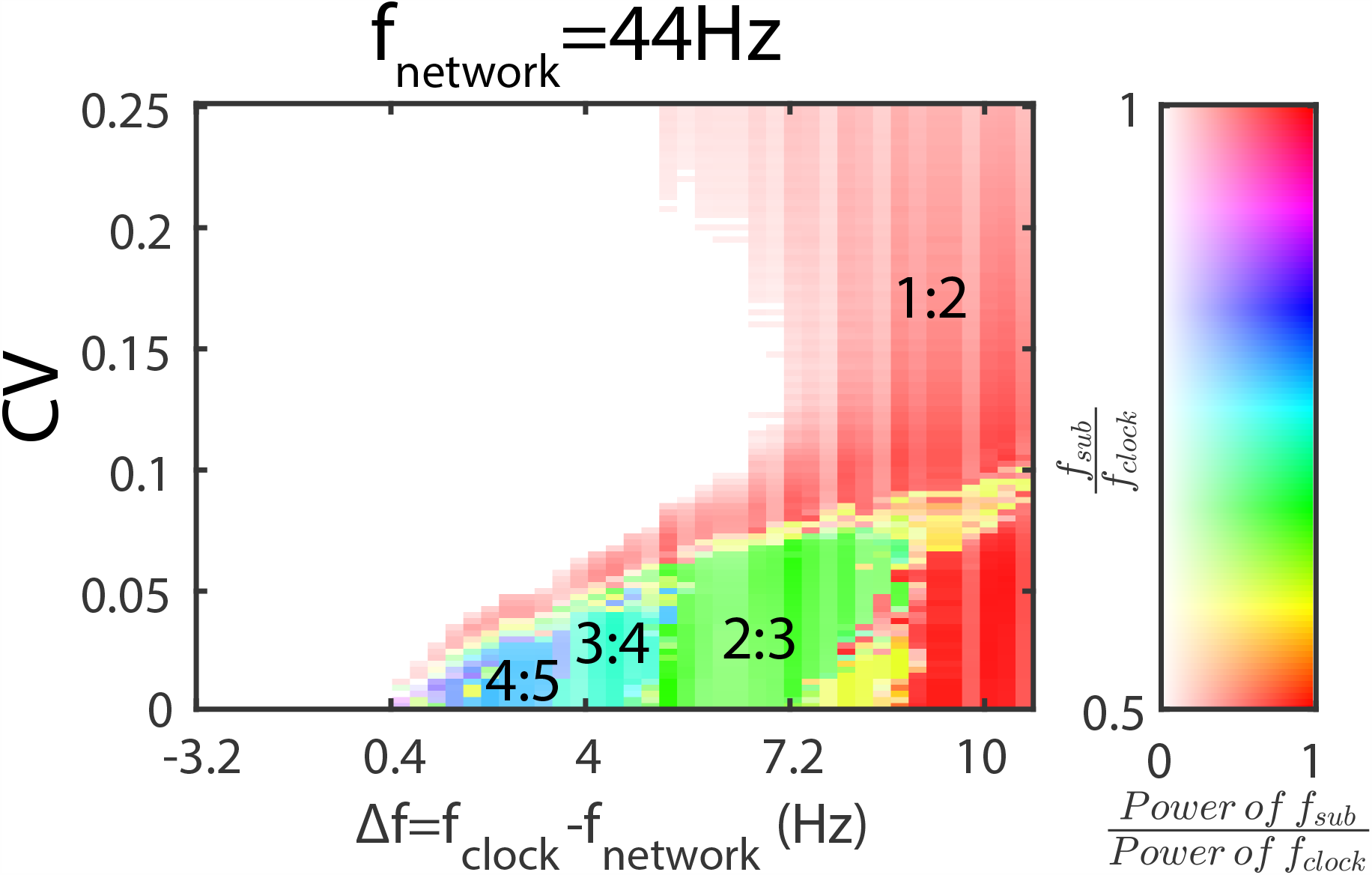
Subharmonic response of the ING rhythm with a longer synaptic delay within the network (*τ*_*d*_ = 5 ms) receiving periodic inhibitory input. For each value of the input heterogeneity, the natural frequency *f*_*network*_ was kept constant (*f*_*network*_ = 44 Hz) by adjusting the mean input strength *I*^(*I*)^. The range of detuning where increasing heterogeneity induced a type 3 synchronization became wider compared to Fig.6B, where *τ*_*d*_ = 3 ms. *W* ^(*ext*)^ = 1.2 × 10^*−*3^.

